# Synergic Renoprotective Effects Of Combined ASC Therapy With RAAS Blockade In Experimental Advanced CKD

**DOI:** 10.1101/2021.09.20.461095

**Authors:** Marina PC Maires, Krislley R Pereira, Everidiene KVB Silva, Victor HR Souza, Flavio Teles, Paulyana F Barbosa, Margoth R Garnica, Felipe M Ornellas, Irene L Noronha, Camilla Fanelli

## Abstract

Global prevalence of chronic kidney disease (CKD) has increased considerably in the recent decades. Overactivity of the renin-angiotensin-aldosterone system (RAAS), associated to renal inflammation and fibrosis contribute to its evolution. The treatments currently employed to control CKD progression are limited and mainly based on the pharmacological inhibition of RAAS, associated with diuretics and immunosuppressive drugs. However, this conservative management promotes only partial deceleration of CKD evolution, and does not completely avoid the progression of the disease and the loss of renal function, which motivates the medical and scientific community to investigate new therapeutic approaches to detain renal inflammation / fibrosis and CKD progression. Recent studies have shown the application of mesenchymal stem cells (mSC) to exert beneficial effects on the renal tissue of animals submitted to experimental models of CKD. In this context, the aim of the present study was to evaluate the effects of subcapsular application of adipose tissue-derived mSC (ASC) in rats submitted to the 5/6 renal ablation model, 15 days after the establishment of CKD, when the nephropathy was already severe. We also verify whether ASC associated to Losartan, would promote greater renoprotection when compared to the respective monotherapies. Animals were followed until 30 days of CKD, when body weight, systolic blood pressure, biochemical, histological, immunohistochemical and gene expression analysis were performed. The combination of ASC and Losartan was more effective than Losartan monotherapy in reducing systolic blood pressure and glomerulosclerosis, and also promoted the complete normalization of proteinuria and albuminuria, a significant reduction in renal interstitial macrophage infiltration and downregulation of renal IL-6 gene expression. The beneficial effects of ACS are possibly due to the immunomodulatory and anti-inflammatory role of factors secreted by these cells, modulating the local immune response. Although studies are still required, our results demonstrated that a subcapsular inoculation of ASC, associated with the administration of Losartan, exerted additional renoprotective effect in rats submitted to a severe model of established CKD, when compared to Losartan monotherapy, thus suggesting ASC may be a potential adjuvant to RAAS-blockade therapy currently employed in the conservative management of CKD.

## INTRODUCTION

The prevalence of chronic kidney disease (CKD) has been increasing worldwide in recent decades [1, 2]. It is estimated that CKD affects around 10% of the global population, leading to progressive loss of renal function and the need for renal replacement therapy (RRT) [3]. Regardless of its etiology, CKD usually progresses with hemodynamic changes, such as increased glomerular pressure and the emergence of systemic hypertension, loss of the kidney filtration barrier selectivity, characterized by proteinuria and albuminuria, serum urea nitrogen and creatinine retention, and the establishment of local renal inflammation, with leukocyte infiltration, increased extracellular matrix (ECM) deposition, development of renal fibrosis and progressive renal failure [4, 5].

Some of the pathophysiological mechanisms known to contribute to CKD progression include the overactivity of the renin-angiotensin-aldosterone system (RAAS), with consequent increase in the biological effects of Angiotensin II (AII), and the activation of local innate and adaptive immune responses which trigger the production of pro-inflammatory mediators, leukocyte recruitment and ECM synthesis [4–9]. Therefore, pharmacological suppression of RAAS with AII receptor blockers (ARBs), such as Losartan, and/or AII converting enzyme (ACE) inhibitors (ACEIs) are the most employed therapeutic strategy in the conservative treatment of CKD. However, although widely applied, RAAS inhibitors alone are not able to revert kidney damage, nor to completely detain CKD progression. Until the present moment there is no specific drug or therapy to treat CKD and to effectively to stop the progression of renal inflammation and fibrosis at once, which motivates the scientific community to investigate experimental and pre-clinical strategies to detain or retard CKD progression [2].

In this context, recent studies have shown promising results with the therapeutic application of mesenchymal stromal cells (mSC) in reducing renal inflammation and fibrosis, in experimental models of CKD [4, 10–12]. According to the current literature, the administration of mSC would not exert its beneficial effects due to direct cell differentiation and tissue replacement, which was previously believed to be the main biological mechanism responsible for the renoprotection achieved with cell therapy, but through paracrine immunomodulatory effects, which may involve the synthesis and secretion of anti-inflammatory cytokines and interleukins [13, 14]. Cavaglieri and collaborators demonstrated that bone marrow-derived mSC (BMSC) protected Wistar rats submitted to the 5/6 renal ablation model of CKD from the development of hypertension, albuminuria and glomerulosclerosis, when administered in the renal subcapsular space, concomitantly with the surgical induction of nephropathy, as a preventive strategy. The authors showed the inoculation via renal subcapsular route to provide effective cell migration through the organ cortex, glomeruli and renal interstitium, presenting satisfactory cell distribution and permanence in the region of interest, throughout the study period [10].

Corroborating these findings, Ornellas and co-authors shown mSC derived from the adipose tissue (ASC) to be effective in preventing renal inflammation, proteinuria and podocyte injury when given to animals submitted to the puromycin aminonucleoside-induced nephrosis model [12]. Xing and collaborators demonstrated that the administration of BMSC through the tail vein of animals submitted to a unilateral ureteral obstruction model exerted a renoprotective effect, attenuating interstitial fibrosis and inhibiting the loss of peritubular capillaries in this model [15]. In line with these findings, Pepineli and co-authors more recently demonstrated that ASC modulates the local inflammatory response in a rat model of chronic allograft nephropathy, induced by the allogeneic kidney transplantation using Fisher 344 rats as donors and Lewis rats as graft recipients. In this elegant model, ASC treatment reduced renal macrophage infiltration, as well as the local expression of proinflammatory and pro-fibrotic factors, such as IL-1β, IL-6, and TNF-α [16].

As mentioned above, there are currently a number of research articles demonstrating positive effects of ASC cell therapy in preventing the development of CKD, in different animal models. However, in none of these studies did the authors verify whether the late renal mSC inoculation would also be effective in halting the progression of the already established nephropathy, or even in reversing kidney damage. Thus, the aim of the present study was to investigate the renoprotective effects of a late single renal subcapsular application of ASC, administered 15 days after the induction of CKD, when all clinical and laboratorial characteristics of renal failure where already evident, in rats submitted to the 5/6 renal ablation model, in order to more closely resemble the clinical settings observed in humans. In addition, we sought to verify whether ASC inoculation, associated to the oral administration of Losartan, would promote greater renoprotection compared to ARB monotherapy, in this same CKD model, in order to analyze whether ASC therapy could be considered as adjuvant to the currently employed pharmacological treatment to control CKD progression.

## MATERIAL AND METHODS

### Animal Model

Ninety male Wistar rats, weighing 220–280 g, were obtained from the local animal facility of the University of São Paulo (USP). Animals were kept at a constant temperature of 23 ± 1°C, 5% relative air humidity, under 12 h artificial light/dark cycle, and had free access to rodent chow and tap water throughout the period of analysis. The experimental protocol followed in the present study was fully approved by the Ethics Committee on the Use of Experimental Animals of the University of São Paulo Medical School (CEUA-FMUSP No 1019/2018).

Seventy of the above-mentioned rats were submitted to the 5/6 renal ablation model of CKD, through a one-step surgical procedure: Animals were anesthetized with inhalation anesthesia with isoflurane (Bio Chimico, Brazil), and underwent a ventral laparotomy, under aseptic conditions. The infarction of two-thirds of the left kidney was obtained by the ligation of two of the three branches of the left renal artery, followed by total nephrectomy of the right kidney. Additional 11 rats, used as controls, were also submitted to isoflurane anesthesia and ventral laparotomy, but with no removal of renal mass (Sham). After surgery, all animals were kept in heated cages until they recovered from anesthesia. Post-operative care included a single dose of antibiotic (IM injection of 0.4 mL/kg Enrofloxacin 5%, Bayer), and 3 doses of analgesic (SC injections of 5 mg/kg Tramadol), one every 24 hours.

### Isolation, Maintenance, Characterization and Inoculation of ASC

Adipose-derived mesenchymal stromal cells (ASC) were isolated from the perigonadal adipose tissue of 3 healthy adult male Wistar rats. The animals were anesthetized by isoflurane inhalation and had their blood directly collected from the abdominal aorta to later obtain normal rat serum (NRS), employed in the *in vitro* experiments. After blood collection, the animals had the perigonadal adipose tissue removed, minced and digested with 0.075% collagenase solution (Sigma-Aldrich, USA). After centrifugation and processing, the cell pellet was resuspended in 10 mL of complete DMEM-Low medium (10% FBS) and plated in cell culture flasks, which were kept in a humid oven at 37°C, 5% CO_2_ (Thermo Fisher Scientific, Marietta, USA). The cells were maintained in culture, being monitored daily under inverted microscopy. Culture medium changes were performed three times a week and cells were trypsinized and replated whenever they reached a confluence between 60% and 80%.

Between the 4^th^ and 6^th^ cell passages, cell viability analysis was performed, using trypan blue staining, followed by the characterization of cell populations through flow cytometry FacsCanto (Becton Dickinson, San Jose, EUA). For this characterization, the presence of specific cellular markers for mSC, such as CD29, CD44, CD90 and CD105, as well as the absence of the pan-leukocyte marker CD45 were verified, using specific monoclonal antibodies [17]. The results were analyzed in the form of histograms of the cell population with positive reaction for each antibody, thus characterizing the mSC population, as shown in **Supplementary Figure 1**. Further cell plasticity tests were performed to verify the ability of ASC to differentiate in osteogenic, chondrogenic and adipogenic cell lines, using a commercially available kit (STEMPRO® Osteocyte / Condrocyte / Adipocyte Differentiation), also shown in **Supplementary Figure 1.** After characterization, ASC were collected from culture flasks and divided into samples containing 2×10^6^ cells resuspended in 10 µL of sterile PBS (cell inoculums). The animals were anesthetized with isoflurane, underwent ventral laparotomy and were submitted to the subcapsular injection of the cell inoculum, followed by the same post-operative care mentioned above.

### Experimental Protocol

The animals had their body weight (BW, g) monitored weekly and their systolic blood pressure (SBP, mmHg) measured with an automated optoelectronic device (Visitech Systems, Apex, NC) every 15 days, when 24-hour urine samples were also collected to verify changes in urinary volume (UV, mL), urinary protein excretion (UPE, mg/24h) by colorimetric analysis (Kit Sensiprot, Labtest # 36, Brazil) and urinary albumin excretion (UAE, mg/24h), by radial immunodiffusion, using a specific anti-rat albumin antibody (MPBiomedicals LLC #55711, USA) [18].

On the 15^th^ day after renal ablation, rats were distributed into five experimental groups, randomized according to their basal UPE, as follows: basal CKD (N=11), euthanized 15 days after CKD induction; CKD (N=13), kept untreated until the 30^th^ day after 5/6 renal ablation; CKD+ASC (N= 15), that received a subcapsular injection of ASC after 15 days of renal ablation and were followed until the 30^th^ day after CKD induction; CKD+LOS (N=12), that received 50 mg/Kg/day of Losartan, diluted in drinking water, from the 15^th^ to the 30^th^ days after CKD induction; and CKD+ASC+LOS (N=14), that received both the ASC subcapsular injection and the oral treatment with Losartan, as illustrated in **Supplementary Figure 2.** Additionally, 3 animals submitted to CKD induction were euthanized to collect total blood and obtain uremic rat serum (URS) for the development of *in vitro* experiments, and the 2 remaining animals submitted to renal ablation were employed for the *in vivo* detection of ASC, as described later on. At the end of the study period, animals were once more anesthetized with isoflurane and submitted to a xipho-pubic laparotomy. The abdominal aorta was punctured, and blood samples were collected to measure serum creatinine (Scr, mg/dL) and blood urea nitrogen (BUN, mg/dL), using commercially available kits (Creatinina #35 Kit and Urea CE # 27 Kit, Labtest, Brazil). The estimated creatinine clearance (CrCl, mg/min) was obtained by measuring the urinary creatinine concentration (Ucr, mg/dL), with the same colorimetric kit, and performing the following calculations: [(Ucr x UV) / Scr] / 1440. Further corrections for rat body surface area (RBSA ≅ 357 cm^2^) were obtained by dividing this result by 0.0357 (CrCl, mg/min/BSA).

The left kidney was removed and weighted. For hypertrophy analysis we performed the following calculation = (left kidney weight / final body weight) x 1000. Kidney samples were then fractionated, half of it was briefly fixed in Du Boscq-Brasil solution for 30 minutes, followed by fixation with buffered paraformaldehyde (pH: 7.4) for 24-72 hours, for further histological and immunohistochemical analysis. The remaining half of the left kidney was quickly frozen in liquid nitrogen and kept at −80⁰C for further analysis of gene expression.

### *In vivo* Detection of ASC

In order to verify the homing and stability of ASC in the renal parenchyma, 2 additional animals, submitted to the 5/6 renal ablation model of CKD were subjected to renal subcapsular injection of 2×10^6^ labeled ASC, after 15 days of CKD induction. Briefly, the nuclear stain 4-6 diamidino-2-phenylindole dihydrochloride (DAPI; Sigma-Aldrich), was added to the culture flasks of ASC in the 4^th^ cell and incubated for 1 hour in a humid oven at 37°C, 5% CO^2^. The cells were harvested, counted and inoculated into the rat renal subcapsular space. One of the animals was euthanized after 24h of cell inoculation, and the other one was kept until 30 days of CKD induction (15 days after ASC injection). The animals were euthanized by IP injection of a lethal dose of Thiopental (0.1 g/rat), kidneys were rapidly frozen and processed in histological sections of 4 μm, which were fixed in acetone, stained with 0.6% Evans blue and analyzed using a fluorescence microscope [10].

### Histological and Immunohistochemical Analysis

The fixed renal fragments were dehydrated, diaphanized and included in paraffin blocks from which 4-μm-thick tissue sections were obtained. For all histological and immunohistochemical analyses, kidney sections were deparaffinized and rehydrated through a sequence of xylol and alcohol baths.

The presence of glomerular architecture disruption was verified in the renal sections of animals from the different experimental groups, stained by the Periodic Acid-Schiff (PAS) technique. The percentage of glomerulosclerosis (GS%) was assessed by the blinded analysis of 50 glomeruli of each animal, under a final magnification of 400x. The extent of renal cortical interstitial fibrosis was estimated in renal sections stained with Masson’s Trichrome. The percentage of interstitial fibrosis was determined by the point-counting technique, in 30 consecutive microscopic fields, under a final magnification of 200x [19].

Immunohistochemistry was employed to detect myofibroblasts, by the presence of the α-smooth muscle actin protein (α-SMA), macrophages (CD68), T-lymphocytes (CD3), ZO1 constitutive glomerular protein, and proliferating cells (PCNA), in the renal samples of experimental animals. After dewaxing, 4 μm thick slices were subjected to microwave heating in pH 6.0 citrate buffer for antigen retrieval. For CD68 and α-SMA, the immunophosphatase technique was performed. The primary monoclonal mouse anti α-SMA (Sigma, #A2547) and monoclonal mouse anti-CD68 antibodies (Serotec, #MCA341R) were used, followed by the secondary biotinylated anti-mouse antibody (Vector # BA2001). Reactions were developed with Fast Red TR salt (Merk, #F6760), as previously described [17]. For CD3, PCNA and ZO1, the immunoperoxidase technique was performed, employing the primary polyclonal rabbit anti-CD3 (DAKO #A4052), monoclonal mouse anti-PCNA (Dako, #M0879) and polyclonal rabbit anti-ZO-1 (ZYMED 617300) antibodies, respectively. Slides were developed with DAB (Dako, Carpinteria, CA, USA) [Machado, 2008]. Quantitative analysis of immunohistochemistry was performed in a blinded fashion. Renal cortical interstitial infiltration by macrophages and T-lymphocytes, as well as cell proliferation were evaluated by counting the number of positive cells for CD68, CD3 and PCNA, in at least 30 microscopic fields for each animal, under a final 400x magnification. The percentage of renal interstitial area occupied by α-SMA was achieved using the same point-counting technique previously described. Finally, structural integrity of the glomerular filtration barrier was verified through the percentage of glomerular area occupied by ZO-1, in at least 25 glomeruli, under a 400x magnification.

### Gene Expression Analysis

Quantitative reverse transcription PCR (RT-qPCR) analyses were performed to assess renal cortical expression of genes coding for pro and anti-inflammatory cytokines, such as: IL-1β, IL-2, IL-4, IL-6 and IL-10. For this purpose, the total RNA of the renal fragments, previously frozen in liquid nitrogen and kept at −80°C, was extracted using Trizol (Ambion Thermo Fischer #15596018). Constitutive gDNA was eliminated from RNA samples using the Turbo DNAse free^TM^ kit (Invitrogen #AM1907). RT was carried out using M-MLV RT enzyme (Promega #M1705) to obtain cDNA. PCR were then conducted using specific pairs of primers (Supplementary Table 1) and the Syber GreenER qPCR Supe Mix Universal (Invitrogen #11762), in the StepOne Plus equipment (Applied Biosystetems - Life Technologies). The *Bact* gene, coding for constitutive Beta-actin protein (BACT), was used as an endogenous control of the reactions and the products of the RT-qPCR reactions were quantified by the ΔΔCt relative method. The primer sequences employed in this technique can be found in Supplementary Table 1.

### *In vitro* Experiments

In order to verify if the treatment with an ARB could somehow impair or reduce the survival of ASC in the renal parenchyma of CKD animals, which would force us to choose another class of RAAS blocker to associate to our experimental cell therapy, we performed *in vitro* experiments to determine if Losartan would be toxic to these cells. At the 4^th^ cell passage, ASC were seeded in cell plates and cultured until achieving 70% of confluence, when the complete medium was removed and replaced with serum-deprived medium for 24h. After this period, cells were finally submitted to the following treatments: 15% of NR, 15% of URS or 15% of URS + 10μM of Losartan, following the literature [20]. Cells were appropriately kept in a humid oven at 37°C, 5% CO_2_ for further 24h and then fixed for immunofluorescence analysis.

### Statistical Analysis

Results were presented as mean ± SE. One-way Analysis of Variance (ANOVA), followed by appropriate Tukey post-test was performed to compare all the groups. All calculations were performed using the GraphPad Prism® software version 7.0, and p values below 0.05 were considered significant [21].

## RESULTS

### ASC inoculation associated to Losartan, reversed hypertension and reduced the mortality of animals with CKD

As expected, animals submitted to the model exhibited significant hypertension after 15 days of CKD induction, when compared to time-paired Sham animals (180±8 vs. 133±3, p<0.05). SBP raised in the untreated CKD group (205±6), as well as in the animals that received only the cell therapy (CKD+ASC; 203±8), which remained significantly higher than that observed in Sham at 30 days of renal ablation (140±2), in both of these groups. Losartan treatment promoted a decrease in SBP, which was statistically different in the group CKD+LOS compared to untreated CKD 30d (174±9 vs. 205±6, p<0.05). Surprisingly, the association of this pharmacological treatment with the subcapsular ASC inoculation promoted an even more pronounced reduction in arterial hypertension in the CKD+ASC+LOS group, in which SBP values were not statistically different from those of Sham (163±9 p>0.05 vs.140±2). The follow-up of SBP in each experimental group, along the study period, can be seen in figure **1A.** To further analyze the impact of the experimental treatments on the evolution of systemic hypertension in the 5/6 model, we calculated the ΔSBP, obtained from the following subtraction: (final SBP, at 30d) – (basal SBP, obtained before the start of treatments, at 15d of CKD induction). This calculation was made individually for each animal, and the mean and standard error values were used in the construction of the graph shown in figure **1B.** As observed, the untreated CKD animals had an increase in SBP of approximately 70% between the 15^th^ and 30^th^ days of analysis, while the monotherapies slowed down this progression: The SBP increase in the CKD+ASC group was around 20% and in CKD+LOS group, nearly only 5%. Remarkably, the ASC+LOS association promoted the reversal of hypertension, as SBP values decreased in CKD+ASC+LOS 30d animals by around 20% after the treatments.

**Figure 1:**
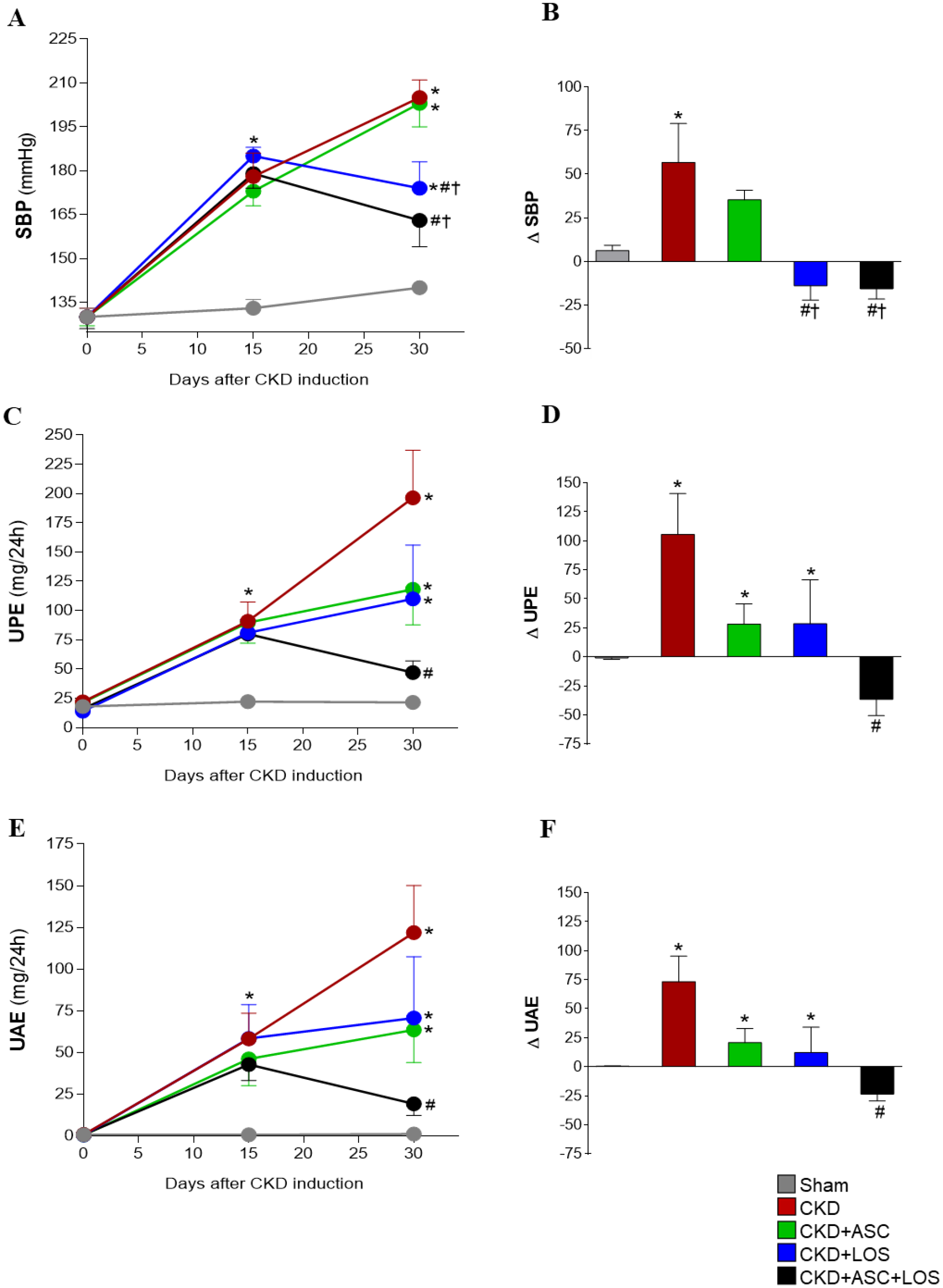
Systolic blood pressure (SBP, mmHg), urinary protein (UPE, mg/24h) and albumin (UAE, mg/24h) excretion of the animals of each experimental group: **(A)** SBP, **(C)** UPE, **(E)** UAE time-course graphs. **(B)** SBP, **(D)** UPE, **(F)** UAE delta bar-graphs, obtained by subtracting the values observed at 30 days from those obtained at 15 days, before the beginning of treatments. Statistical differences are: *p<0.05 vs. Sham, #p<0.05 vs. CKD, †p<0.05 vs. CKD+ASC, ‡p<0.05 vs. CKD+LOS, in the respective time points.

Accordingly, the percentage of survival among the animals between the 15^th^ and the 30^th^ day after CKD induction followed the dynamics of hypertension in this model. The survival rate was around 90% in the groups that did not receive Losartan (untreated CKD and CKD+ASC groups), while there was no mortality in the groups CKD+LOS and CKD+ASC+LOS in this same period (100% of survival), as shown in **Supplementary Table 2.**

### The association of ASC to Losartan promoted the regression of proteinuria and albuminuria in the established CKD model

As can be seen in Figure **1C** and **1E**, the baseline protein and albumin urinary excretion values of the animals at time zero, before CKD induction, were compatible with physiological levels of normal renal function, and very similar among all the animals included in the study. At 15 days after renal ablation, the animals already exhibited significant proteinuria and albuminuria, when compared to Sham (81±20 vs. 20±2, and 51±19 vs. 1±1, p<0.05). At this point, rats underwent to the remnant model were subdivided into CKD (untreated), CKD+ASC, CKD+LOS and CKD+ASC+LOS groups, based on similar initial proteinuria and albuminuria values. In untreated CKD animals, both proteinuria (196±41) and albuminuria (122±28) increased significantly between day 15 and day 30 after 5/6 renal ablation, reaching approximately twice the value observed at 15 days, at the end of the study (Figure **1D** and **1F**). The administration of both ASC or LOS monotherapies promoted deceleration of the progression of proteinuria and albuminuria in CKD+ASC (118±30 and 63±20) and CKD+LOS (110±46 and 71±37), respectively. Surprisingly, CKD+ASC+LOS animals exhibited significant regression of both proteinuria (47±10) and albuminuria (19±7), at 30 days after renal ablation. In this group the final values of protein and albumin urinary excretion were not statistically different from those observed in the Sham group.

### Detection of ASC in the renal parenchyma of CKD rats after 15 days of renal subcapsular inoculation

As illustrated in the upper part of figure 2, 15d after CKD induction, 2 animals received a subcapsular injection of 2×10^6^ ASC, previously labeled with DAPI (**2A**). One of these rats was euthanized after 24h of ASC inoculation, and the other one was kept for more 15 days after cell therapy. Evans blue-stained renal slides were analyzed under fluorescence microscope, as represented in the illustrative microphotographs in **2B**. As shown, DAPI-labeled ASC were easily detected in the renal parenchyma of CKD animals after 24h and 15d of subcapsular inoculation (fluorescent blue cells).

**Figure 2:**
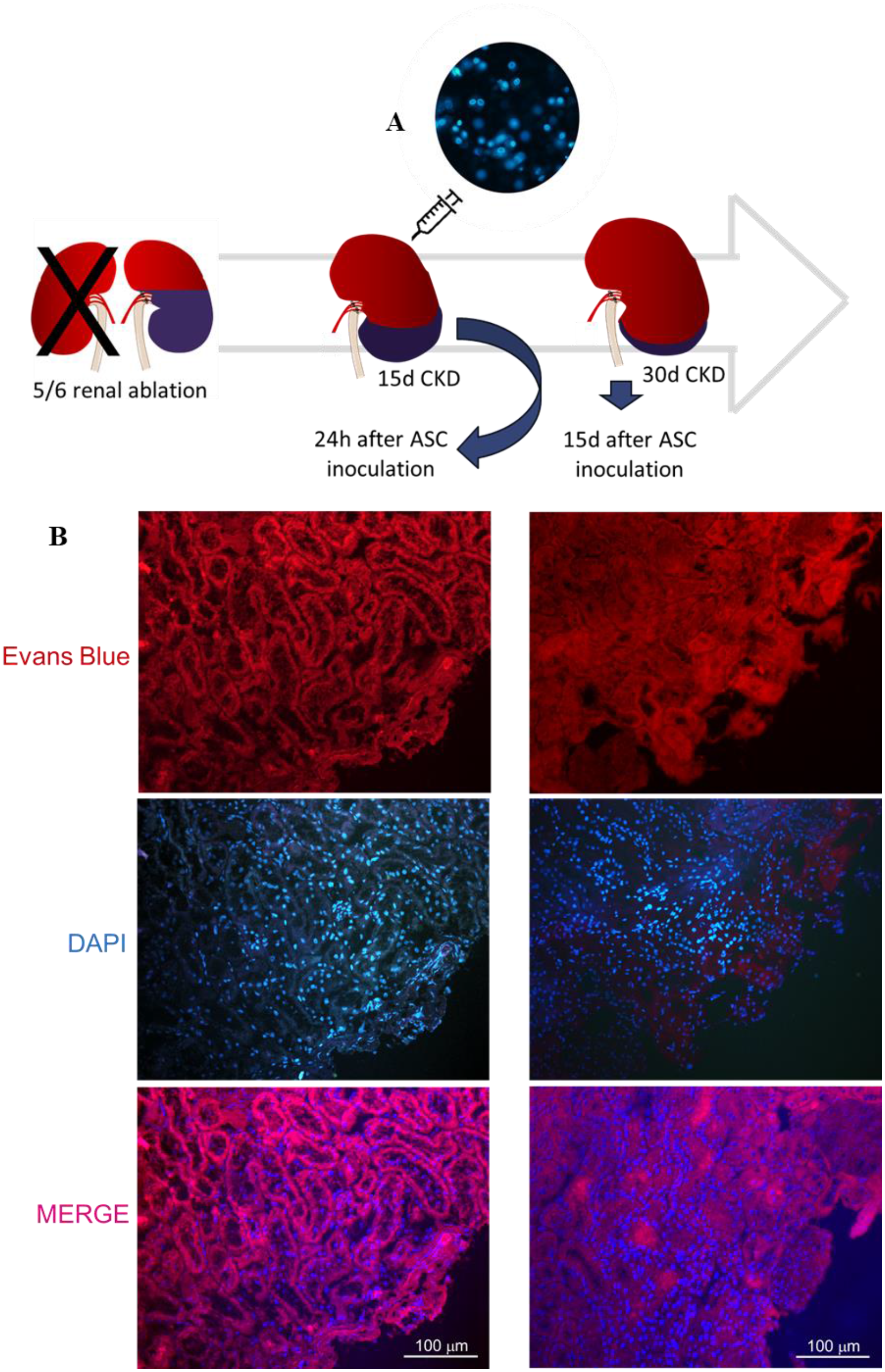
In vivo detection of ASC: After 15d of CKD induction, animals received a subcapsular injection of previously DAPI-labeled ASC **(A)**, which could be detected 24h and also 15d after inoculation, under a final 200x magnification, using a fluorescence microscope **(B)**.

### ASC inoculation combined to Losartan treatment averted the progression of structural glomerular damage in experimental CKD

Glomerular architecture was evaluated in PAS-stained renal sections, under a final 400x magnification, as illustrated in figure **3A**. The percentage of glomerulosclerosis (GS%) in each experimental group was represented as a bar graph, shown in figure **3B**. Corroborating previous findings, animals submitted to the 5/6 renal ablation model already presented a numerically high GS% after 15 days of CKD induction (21±5), which markedly progressed over time, achieving statistically significant values at 30 days of CKD in both untreated and CKD+LOS groups, when compared to time-paired Sham rats (28±6 and 23±6 vs. 3±1, p<0.05, respectively). ASC inoculation alone (17±5) or in association to Losartan (13±2) significantly prevented GS progression in CKD+ASC and CKD+ASC+LOS groups, which did not differ statistically from control group.

**Figure 3:**
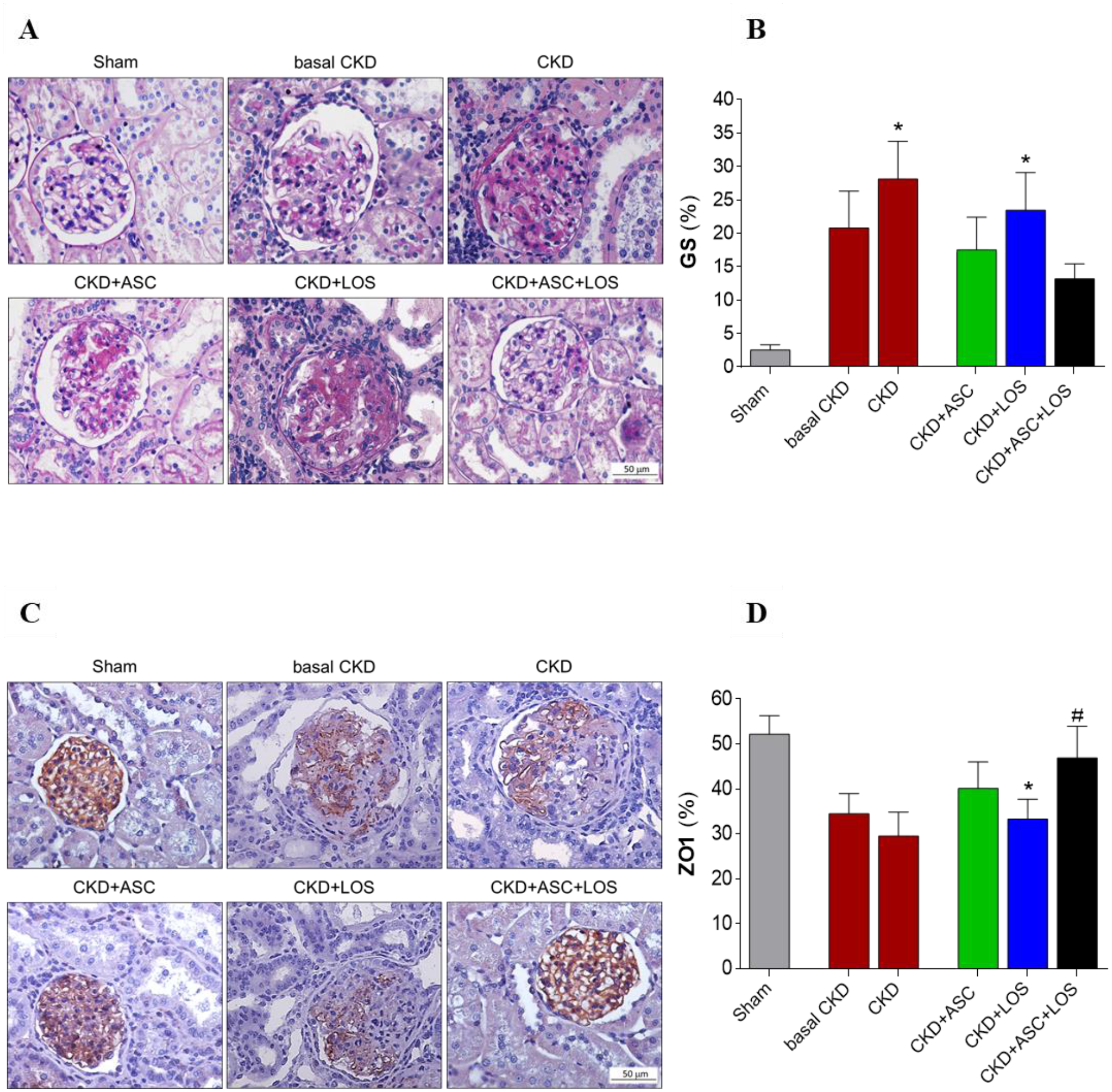
Glomerular architecture: **(A)** Illustrative microphotographs of PAS stained sections of the animals of each experimental group, at the end of the study. **(B)** Bar graphs of the percentage of glomerulosclerosis (GS) in the animals included in the protocol. **(C)** Illustrative microphotographs of immunohistochemistry for ZO1 in each experimental group. **(D)** Bar graphs of the percentage of glomerular area occupied by ZO1 in the animals included in each experimental group, at the end of the study. Statistical differences are: *p<0.05 vs. Sham, §p<0.05 vs. basal CKD, #p<0.05 vs. CKD, †p<0.05 vs. CKD+ASC, ‡p<0.05 vs. CKD+LOS.

We further analyzed the percentage of glomerular area occupied by the constitutive protein ZO1, related to the integrity of the glomerular filtration barrier. Illustrative microphotographs of renal sections submitted to immunohistochemistry for ZO1 can be seen in figure 3C, ZO1% bar graphs, in figure 3D. All CKD animals exhibited reduced percentage of the glomerular area occupied by ZO1, when compared to Sham; except for the rats treated with ASC+LOS association, in which ZO1% was statistically similar to that observed in Sham animals (53±3 vs. 52±4 p>0.05).

### Subcapsular ASC inoculation was as effective as Losartan in attenuating the renal fibrosis in the remnant kidney model

Renal cortical interstitial fibrosis was evaluated in Masson’s trichrome sections by the presence of interstitial collagen, stained in blue, under a final 200x magnification, as illustrated in figure **4A**. Moreover, immunohistochemistry for α-SMA was employed to access the presence of renal interstitial myofibroblasts, major effector cells of fibrogenesis, stained in red, under a final 200x magnification, as shown in figure **4C**. The percentage of interstitial fibrosis (INT%) and renal interstitial area occupied by α-SMA, in each experimental group, were represented as bar graphs, shown in figures **4B** and **4D**, respectively. At 15 days of renal ablation, basal CKD animals already showed a significant increase in the renal cortical interstitial area occupied by collagen, as well as in the fraction of the cortical interstitium occupied by α-SMA, when compared to the control group (2.0±0.3 and 3.6±0.7 vs. 1.1±0.2 and 1.0±0.1, p<0.05), characterizing the presence of interstitial fibrosis, which persisted after 30 days of CKD induction in untreated rats (5.7±1.5 and 3.5±0.5 vs. 1.1±0.2 and 1.0±0.1, p<0.05). Animals treated between 15 and 30^h^ days of CKD with LOS or ASC monotherapies or with ASC+LOS association, showed a numerical reduction of both INT% and interstitial α-SMA which was equivalent among all treated groups.

**Figure 4:**
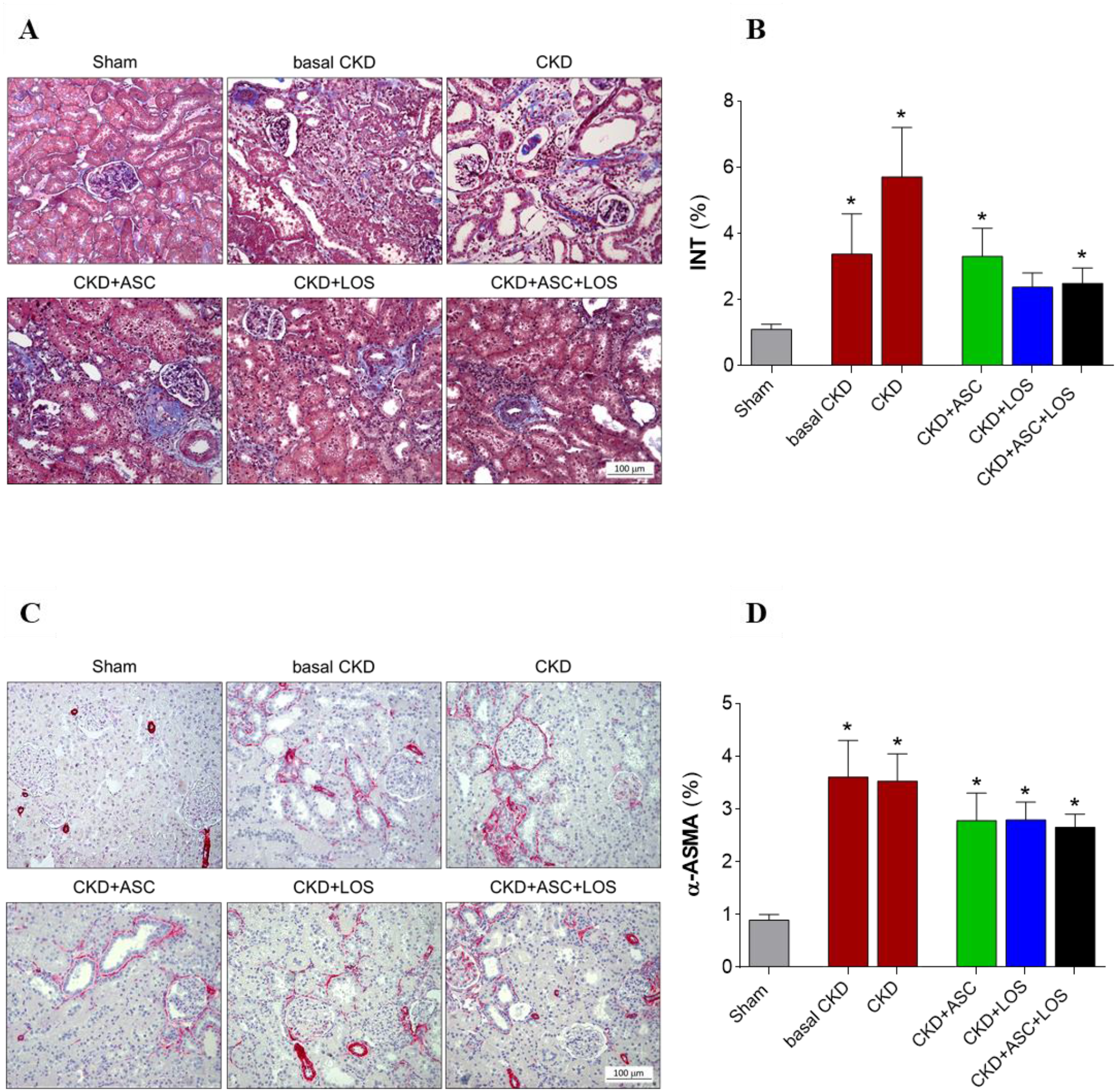
Interstitial fibrosis and myofibroblasts infiltration. **(A)** Illustrative microphotographs of Masson’s Trichrome staining in the experimental groups. **(B)** Bar graphs representing the percentage of the renal interstitial area occupied by fibrosis in the studied animals. **(C)** Microphotographs illustrating the immunohistochemistry for α-ASMA to determine the presence of interstitial myofibroblasts in the renal parenchyma of the studied animals. **(D)** Bar graphs representing the percentage of interstitial area occupied by α-ASMA. Statistical differences are: *p<0.05 vs. Sham, §p<0.05 vs. basal CKD, #p<0.05 vs. CKD, †p<0.05 vs. CKD+ASC, ‡p<0.05 vs. CKD+LOS.

### The association of ASC to Losartan interrupted the progression of renal cortical inflammation and downregulated IL-6 gene overexpression in the 5/6 ablation model of CKD

Interstitial inflammation is one of the main features of CKD progression. Here we evaluated this process by both the presence and intensity of renal infiltration by leukocytes and by the increased proliferation of interstitial cells in the renal parenchyma. Illustrative microphotographs of immunohistochemistry for CD68+ Macrophages (**5A**), CD3+ T-Lymphocytes (**5C**) and PCNA+ interstitial cells (**5E**) are presented in figure 5. As can be seen in the bar graphs in figures **5B**, **5D** and **5F**, after 15 days of CKD induction, animals already presented marked macrophage infiltration and significant increase in the presence of T-lymphocytes and PCNA+ interstitial cells, when compared to the control animals (93±18, 53±10 and 101±16 vs. 36±7, 14±3 and 24±3, respectively). Renal inflammation progressed over time in untreated rats, reaching statistically significant values after 30 days of CKD in all analyzed parameters: (153±35 for CD68, 132±30 for CD3 and 135±27 for PCNA vs. 36±7, 14±3 and 24±3, respectively, p<0.05). ASC or Losartan monotherapies were not able to detain the progression of renal inflammation in the ablation model. However, the association of ASC+LOS promoted significant reduction of macrophage infiltration (56±7 vs. 153±35 p<0.05), and a relevant reduction in both interstitial lymphocytes (76±12 vs. 132±30) and PCNA+ cells (64±14 vs. 135±27). The results of our RT-qPCR analyses of the local renal expression of some of the main pro and anti-inflammatory genes related to CKD progression can be seen as bar graphs in figure 6. After 15 days of CKD induction, rats already exhibited IL-1β, IL-2, IL-4, IL-6, IL-10 and TGF-β overexpression, which remained up to 30 days of renal ablation. It is noteworthy that the association of ASC to Losartan promoted further upregulation of IL-4 (6C) and significant downregulation of IL-6 (6D) gene expressions.

**Figure 5:**
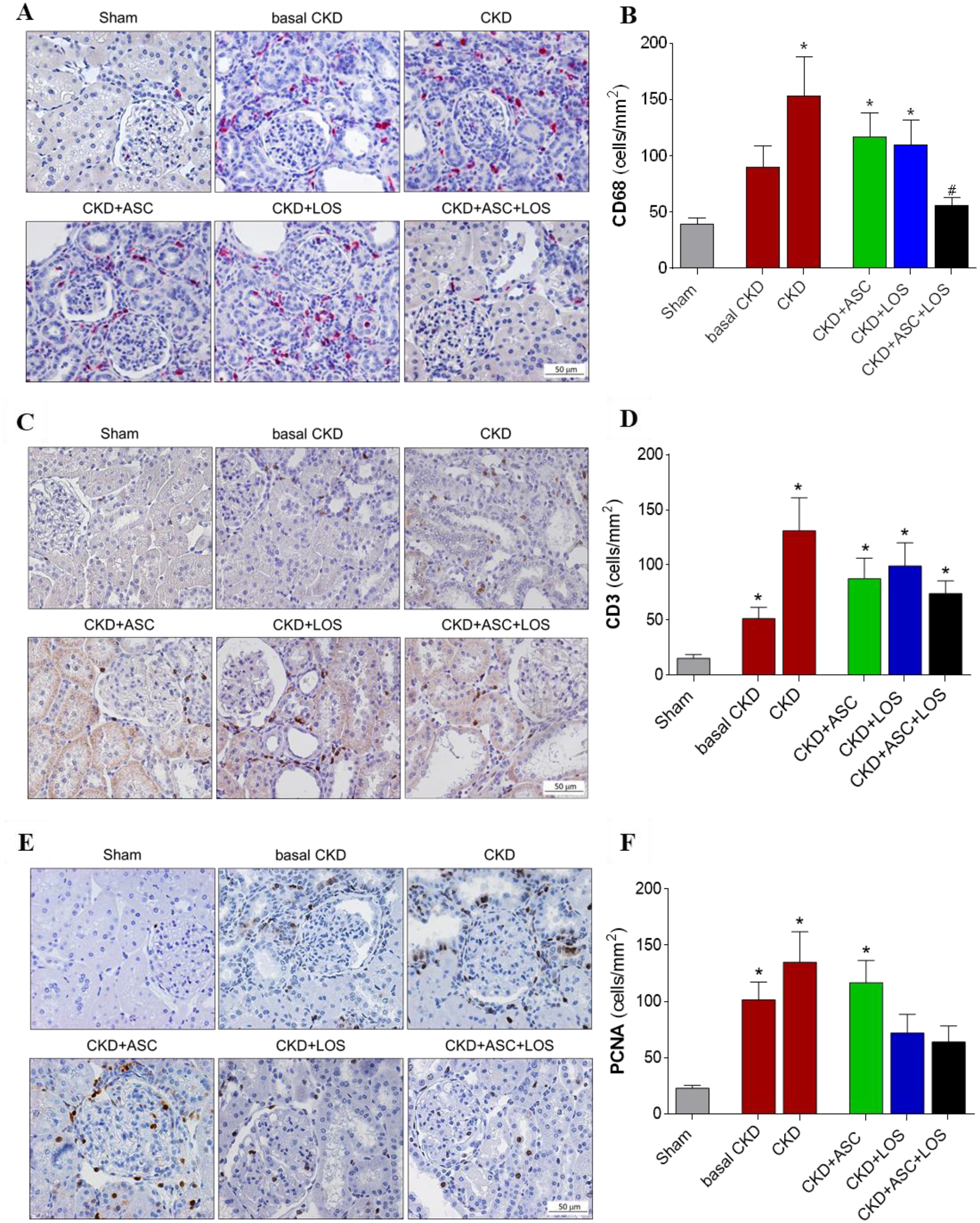
Local renal inflammation: Illustrative microphotographs of immunohistochemistry for **(A)** macrophages (CD68), **(C)** T-cells (CD3) and **(E)** Proliferating interstitial cells (PCNA), in each experimental group. Bar graphs representing the number of **(B)** macrophages, **(D)** T-lymphocytes and (F) PCNA (cells/mm2) in the renal cortical interstitial area. Statistical differences are: *p<0.05 vs. Sham, §p<0.05 vs. basal CKD, #p<0.05 vs. CKD, †p<0.05 vs. CKD+ASC, ‡p<0.05 vs. CKD+LOS.

**Figure 6:**
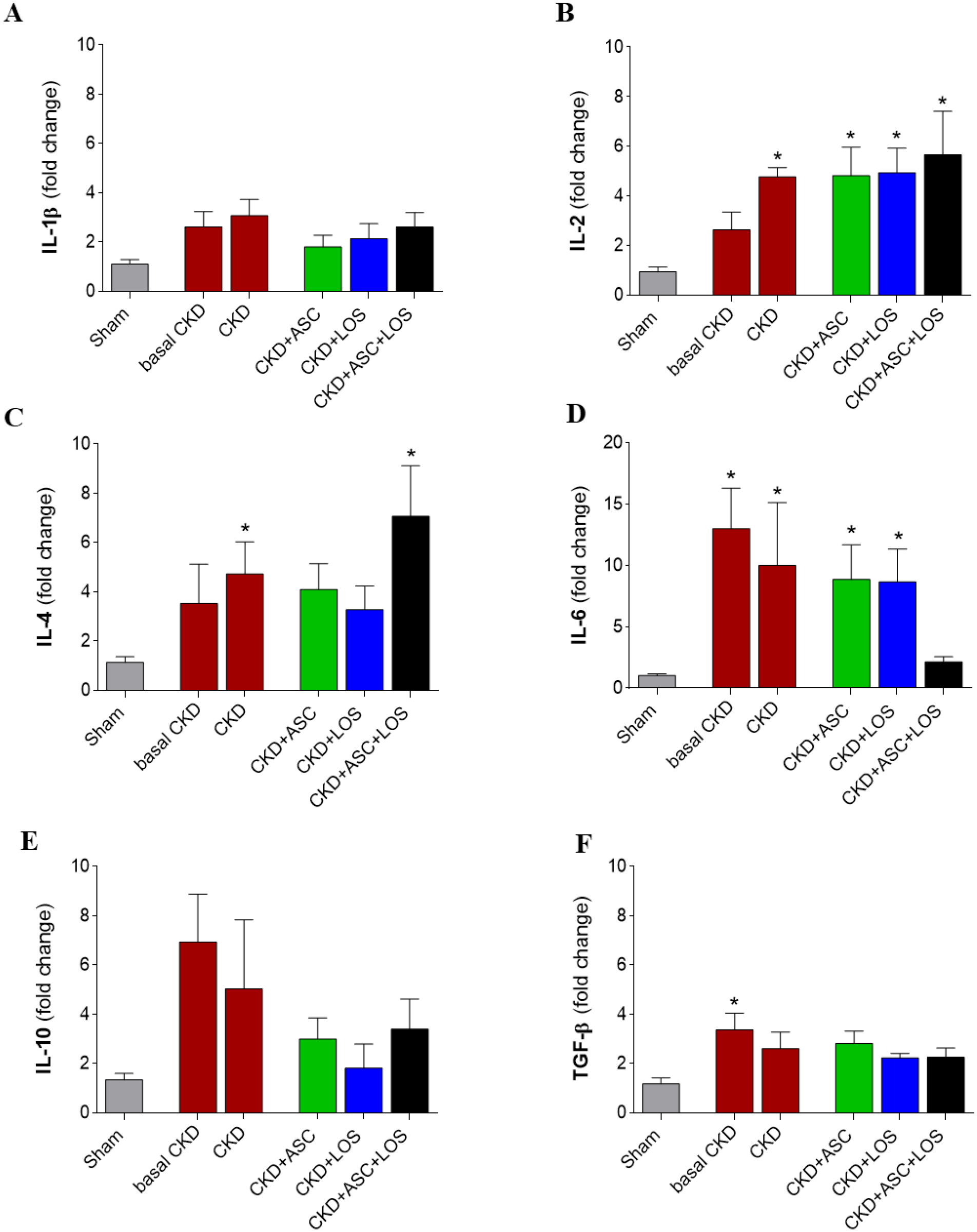
Bar graphs depicting the gene expression results of the following pro-inflammatory and anti-inflammatory genes using the RT qPCR technique: **(A)** IL-1β, **(B)** IL-2, **(C)** IL-4, **(D)** IL-6, **(E)** IL-10 and **(F)** TGF-β. Statistical differences are: *p<0.05 vs. Sham, §p<0.05 vs. basal CKD, #p<0.05 vs. CKD, †p<0.05 vs. CKD+ASC, ‡p<0.05 vs. CKD+LOS.

### Losartan improved cell viability and proliferation of ASC *in vitro*

As illustrated in **Supplementary Figure 3**, cultured ASC stimulated with Losartan *in vitro* exhibited a significant increase in the total number of living cells/mm^2^, as well as in the number of proliferating ASC, which were positive for PCNA in immunofluorescence analysis.

## DISCUSSION

CKD is still a major global health issue. According to recent surveys, by the end of 2020 there were at least 2.5 million of patients receiving renal replacement therapy worldwide, and this number is expected to double by 2030. The high mortality rates and the highly debilitating character of CKD, together with the current lack of an effective treatment to halt its progression, urge the world to seek for new therapeutic strategies to detain the advance of CKD [22, 23]. In the present study we sought to verify the potential renoprotective effects of cell therapy, using a high number of ASC (2 x 10^6^), administered directly in the renal subcapsular area, in Wistar rats submitted to the 5/6 renal ablation model of CKD. In addition, we aimed to investigate the effects of the association of this innovative therapeutic strategy with the pharmacological blockade of RAAS, currently employed in the conservative management of CKD.

As widely described in the literature, the 5/6 renal ablation is a very consistent experimental model to mimic human progressive nephropathy [9,24,25]. Accordingly, in the present study, the animals submitted to this procedure exhibited severe arterial hypertension, loss of selectivity of the glomerular filtration barrier, evidenced by massive proteinuria and albuminuria, along with histological alterations of the renal parenchyma, such as glomerulosclerosis and tubulointerstitial inflammation and fibrosis, as early as 15 days after CKD induction. Similar to what is usually observed in human chronic nephropathy, these parameters worsened over time in untreated CKD animals, reaching exuberant values, and leading to elevated mortality rates after 30 days of renal ablation. Since most of the characteristic features of CKD were completely established 15 days after 5/6 ablation, we choose this point to start the experimental treatments, to better investigate the potential effects achieved by these therapies on patients already affected by CKD.

Our research group choose the renal subcapsular space to inoculate ASC, based on the traditional results obtained by Melgren and collaborators, using pancreatic islets. These authors demonstrated that the renal subcapsular space of diabetic mice offered better conditions for the growth of transplanted pancreatic islets, when compared to other sites of inoculation, such as the liver and spleen [26]. More recently, in an experimental study with rats, Svensson et. al also showed that transplanted pancreatic islets inserted in the renal subcapsular space presented greater revascularization and oxygenation, when compared to islets inoculated directly in the liver of these animals [27]. Corroborating the results obtained by Cavaglieri and collaborators, in an elegant study employing the 5/6 renal ablation model of CKD, our analysis of cell migration demonstrated the effective displacement and distribution of cells through the renal cortex of the animals that received the subcapsular renal inoculation of ASC, which were still present in the renal parenchyma until the end of the study, 30 days after renal ablation, thus consolidating the renal subcapsular pathway as a viable alternative for cell therapy, at least in experimental studies [10].

Here, we demonstrated that ASC alone slowed down the progression of proteinuria, albuminuria and glomerulosclerosis in animals that already exhibited significant renal injury. Furthermore, it partially detained ZO1 depletion observed in animals with CKD, thus preserving the tight junctions between podocyte processes and reducing the loss of macromolecules to the space of Bowman, suggesting a specific protective effect of ASC on the glomeruli. Our findings are in agreement with those of Ornellas et al., which demonstrated BMSC to protect rats submitted to a model of puromycin-induced glomerular injury from podocyte loss, from the effacement of the podocyte processes and from the loss of constitutive glomerular filtration barrier compounds such as nephrin, podocin, synaptopodin and podocalyxin [12]. Additionally, we showed that a single administration of ASC at 15 days of CKD was efficient in detaining the progression of established tubulointerstitial fibrosis, keeping the percentage of interstitial area of CKD+ASC animals at 30 days of renal ablation numerically similar to that observed before ASC injection. Akan and collaborators have recently shown similar results with the intravenous injection of human amnion-derived mSC (hAMSC) in this same CKD model. The authors observed a reduction in the interstitial expression of collagen and TGF-β1 in the renal parenchyma of animals submitted to the cell therapy, accompanied by the increased expression of Bone Morphogenetic Protein-7 (BMP7), a member of the TGF-β superfamily, which counteracts the biological functions of TGF-β1, thus exhibiting anti-fibrotic properties [28, 29]. Therefore, Tang et al. demonstrated that BMSC inoculation promoted a decrease in the renal expression of α-SMA, collagen types I, II, III and TGF-β1, in animals submitted to the adenine-overload model of interstitial fibrosis [30].

The development of RAAS blockers, more than 2 decades ago, was a major breakthrough for the management of CKD until the present days. Pharmacological RAAS blockade reduced the mortality of CKD patients by cardiovascular events, and considerably improved life quality, mainly due to its antihypertensive and antiproteinuric effects. Nevertheless, the reduction in proteinuria obtained with ACEIs or ARBs is insufficient for a number of patients, moreover, these treatments delay but do not prevent CKD progression and renal function deterioration [31, 32]. Corroborating what has been observed in human nephropathy, and in accordance with previous studies [24], here we demonstrated that oral treatment with 50 mg/Kg/d of Losartan between the 15^th^ and the 30^th^ days completely prevented the mortality of rats submitted to the 5/6 renal ablation model. Moreover, RAAS blockade in monotherapy reversed hypertension and slowed down but did not stop the progression of proteinuria and albuminuria.

A number of *in vitro* studies have demonstrated AII to stimulate the proliferation and activation of cultured fibroblasts [33]. Additionally, AII can be described as a growth factor known to play a crucial role in epithelial-to-mesenchymal transdifferentiation in the development of tumor metastasis development, as well as in renal, alveolar and peritoneal epithelial cells [34]. In accordance with the publications employing cultured cells, experimental studies with different CKD models endorse the profibrotic and proinflammatory biological activities of AII, which is described to act as a cytokine, positively regulating renal and immunological cell response of the beneficial effects of RAAS blockade on renal inflammation and fibrosis [33–36]. Accordingly, here we demonstrated that CKD+LOS animals exhibited significantly less renal interstitial fibrosis, and cell proliferation, as well as a slight reduction in both macrophage and lymphocyte infiltration in the renal parenchyma, compared to the untreated CKD rats.

Since the currently employed RAAS blockade is not enough to stop CKD progression, here we associated cell therapy to this oral treatment in order to verify if additional renoprotection would be achieved. This was the first study to associate the injection of ASC with a clinically employed pharmacological approach of RAAS blockade. We chose the ARB Losartan, instead of an ACEI, based on our *in vitro* findings, which showed that cultured ASC, stimulated with uremic rat serum, in order to better mimic the microenvironment of renal subcapsular space of CKD animals, presented greater proliferative capacity when treated with LOS diluted in the culture media, compared to cells cultivated in the absence of this drug, thus proving evidence that RAAS blockage did not impair ASC growth and survival, and even exerted stimulatory effects on these cells [20].

When associated to Losartan treatment, a single subcapsular inoculation of 2×10^6^ ASC promoted a significant improvement in several parameters associated with the progression of CKD in the 5/6 renal ablation model; especially proteinuria and albuminuria, whose values did not only stop progressing, but also regressed with the combined therapy. Surprisingly, ASC+LOS association virtually normalized UPE and UAE in the 5/6 nephrectomy CKD model, since the CKD+ASC+LOS group did not differ from the Sham group in these parameters. Moreover, the association of ASC to LOS promoted the regression of structural glomerular damage by reversing glomerulosclerosis and preserving ZO1 glomerular expression in the CKD animals more efficiently than the monotherapies. We also observed that ASC+LOS significantly detained hypertension and renal interstitial macrophage infiltration more effectively than ASC or LOS alone, also normalizing the renal expression of IL-6, a potent pro inflammatory mediator.

## CONCLUSION

In the present study, we demonstrated for the first time that the association of a subcapsular injection of ASC with RAAS blockade promoted greater renoprotection when compared to both strategies in monotherapy, thus suggesting cellular therapy with ASC to be a potential adjuvant to the current pharmacological approach used for CKD conservative management, although more studies are still required before the clinical use of ASC can be established.

## DATA AVAILABILITY

Full rough data supporting the findings of this study are available from the corresponding author upon request.

## ACKNOWLEDGMENTS

We would like to thank our laboratory team, namely the colleagues: Andreza Aparecida Santos, Anne Carolina da Silva Constantino, Arizla Benicio Vieira de Menezes, Débora Cristina de Sousa, Kelly Ribeiro Cascimiro, Marcia Ribalta and Wellisson Farias Pereira, for their excellent technical support.

## PREPRINT

In accordance with Hindawi politics, the authors are going to deposit the present manuscript in the preprint server medRxiv within the next few days.

## SUPPLEMENTARY MATERIAL

**Supplementary Figure 1:**
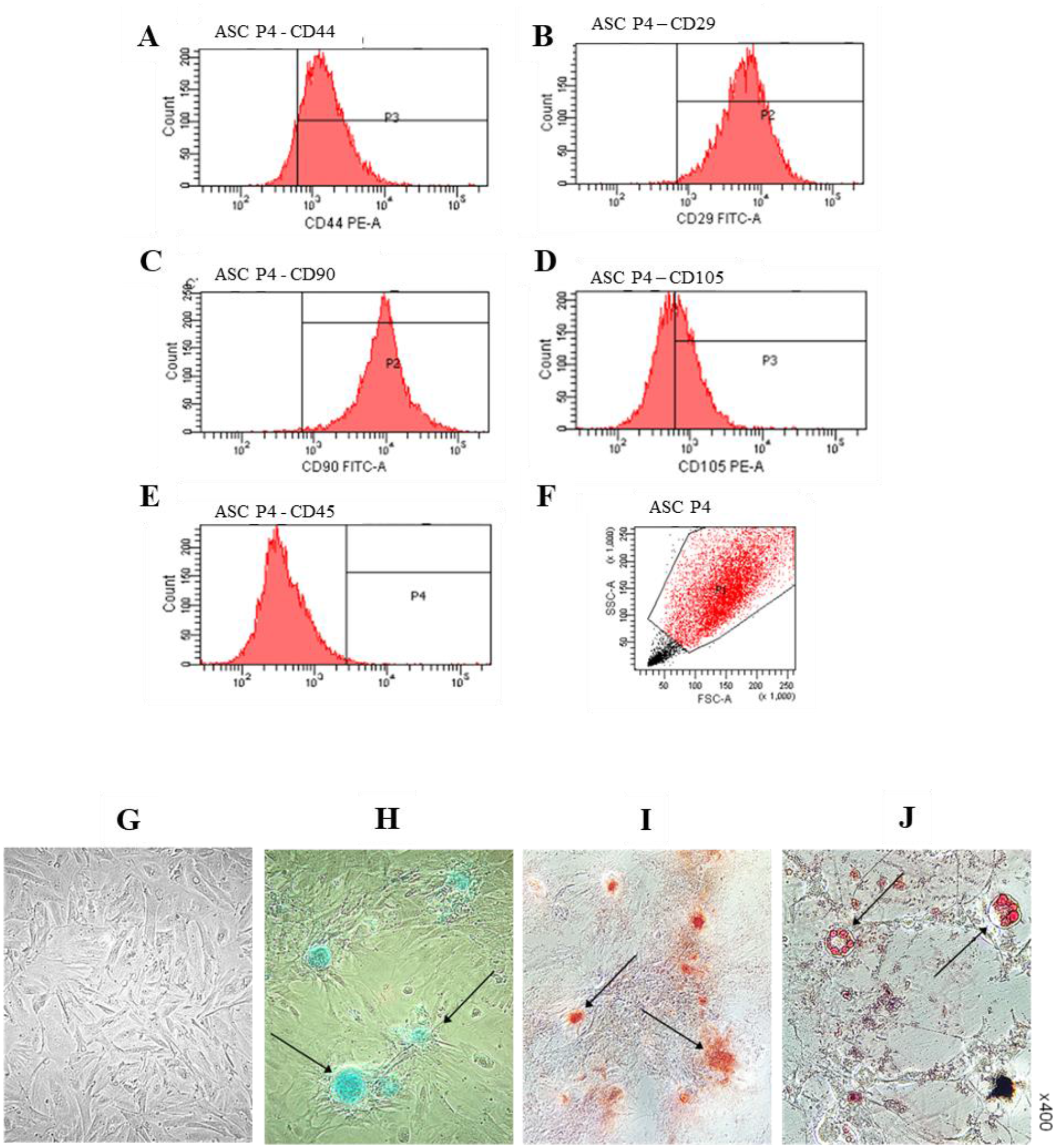
Characterization of ASC at the 4th passage was performed by flow cytometry and cell plasticity tests: Cell population was positive for typical mSC biomarkers: CD44 **(A)**, CD29 **(B)**, CD90 **(C)** and CD105 **(D)**, and negative for the pan leukocyte marker CD45 **(E)**. 10,000 events were analyzed, of which 8,898 were viable **(F)**. ASC phenotype before the plasticity tests is shown in **(G)**. Chondrogenic differentiation was evidenced by the presence of sulfated matrix proteoglycans, highlighted in turquoise blue and stained with Alcian blue **(H)**; the osteogenic differentiation was verified by means of reddish calcium crystals stained with Alizarin Red **(I)** and finally, the adipogenic differentiation was verified by the presence of lipid droplets stained in red by Oil-red O **(J)**. Microphotographs were taken under 400x magnification.

**Supplementary Figure 2:**
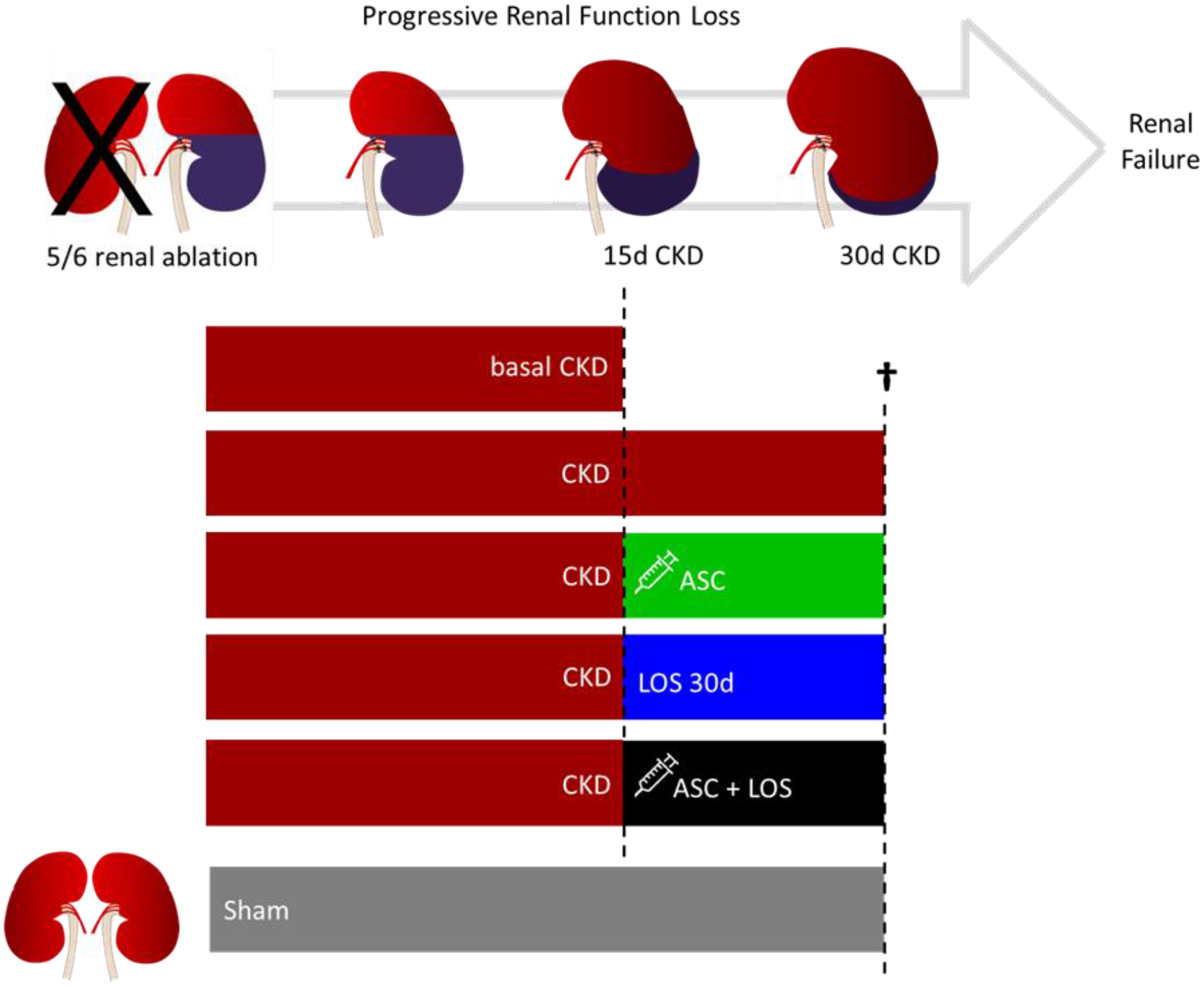
Experimental Protocol. On the 15th day after renal ablation, animals submitted to CKD induction were distributed into five experimental groups, randomized according to their basal UPE, as follows: basal CKD (N=11), euthanized 15 days after 5/6 renal ablation, CKD (N=13), kept untreated for 30 days, CKD+ASC (N=15), that received a subcapsular injection of ASC at the 15th day of CKD and were followed until the 30th day after CKD induction, CKD+LOS (N=12), that received 50 mg/Kg/day of Losartan, diluted in drinking water, from the 15^th^ to the 30^th^ day after renal ablation, and CKD+ASC+LOS (N=14), that received both the ASC subcapsular injection and the oral treatment with Losartan. Sham-operated animals (N=11) were used as control.

**Supplementary Figure 3:**
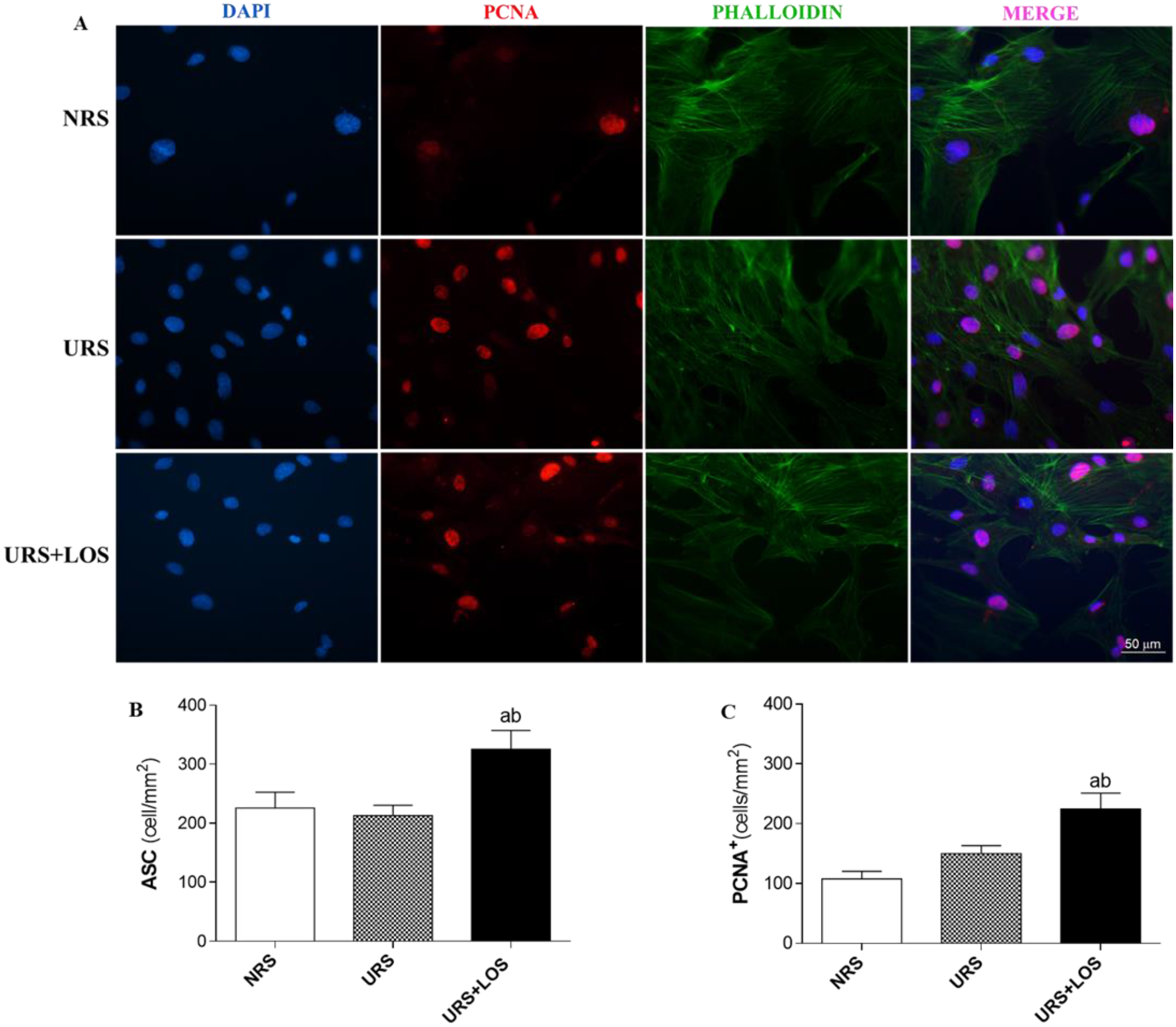
*In vitro* Experiments. **(A)** Illustrative microphotographs of immunofluorescence of cultured ASC, previously treated with: Normal rat serum (NRS), uremic rat serum (URS) or URS+10μM of Losartan (URS+LOS). Cells positive for PCNA are stained in red. The total number of ASC/mm^2^ in each experimental treatment is represented in the bar graph in **B**, while the number of PCNA+ASC/mm^2^ is represented in **C.**

**Supplementary Table 1:**
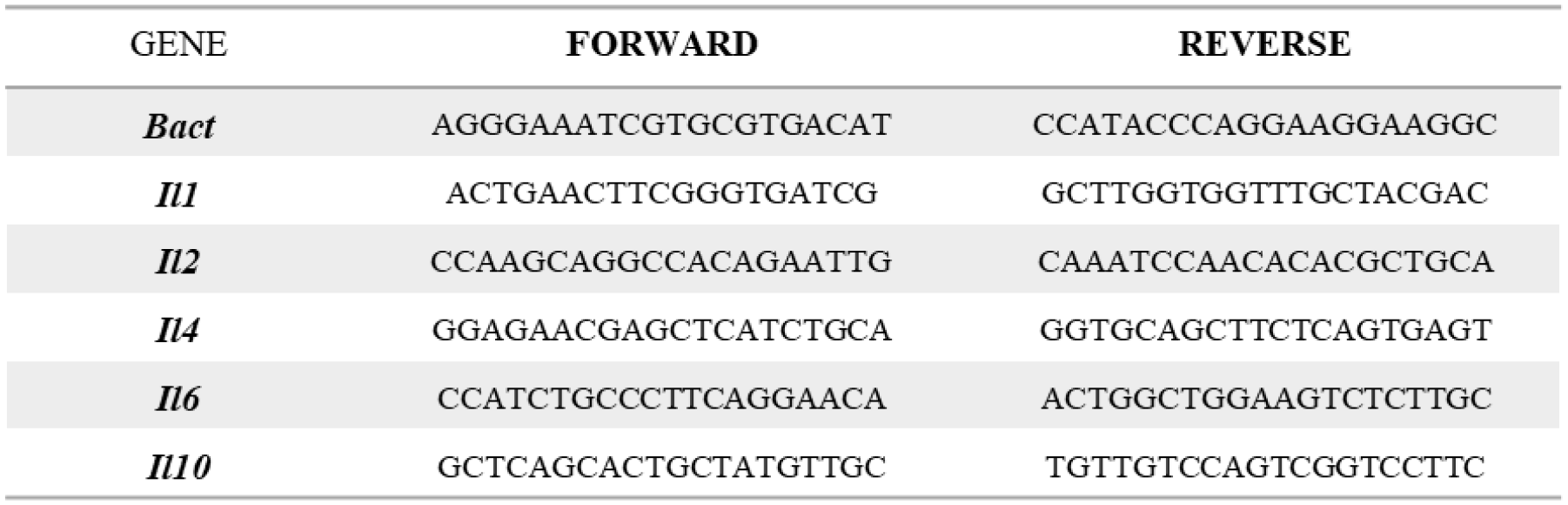
Genes and primer sequences. The housekeeping beta-actin (*Bact*) gene was used as an endogenous control of the PCR reaction.

**Supplementary Table 2:** Final parameters analyzed at the end of the study period: Survival rate (SR; %), Body weight (BW; g), Urinary volume (UV; mL), Serum creatinine concentration (Scr; mg/dL), Urinary creatinine concentration (Ucr; mg/dL), Creatinine clearance (CrCl; mg/min/RBSA), Blood urea nitrogen (BUN; mg/dL) and Renal hypertrophy. *: p<0.05 vs. Sham 30d; §: p<0.05 vs. Basal CKD, #: p<0.05 vs. CKD; †: p<0.05 vs. CKD+ASC; ‡: p<0.05 vs. CKD+LOS.

|  | **Sham 30d**<br>(N=11) | **basal CKD**<br>(N=11) | **CKD**<br>(N=15) | **CKD+ASC**<br>(N=15) | **CKD+LOS**<br>(N=12) | **CKD+ASC+LOS**<br>(N=14) |
| --- | --- | --- | --- | --- | --- | --- |
| **Survival Rate (%)** | 100% | - | 92% | 93% | 100% | 100% |
| **BW (g)** | 356 ± 12 | 276 ± 9* | 304 ± 6* | 286 ± 11* | 289 ± 9* | 296 ± 5* |
| **UV (mL)** | 19 ± 2 | 38 ± 39* | 43±4* | 39±4* | 42±5* | 38±3* |
| **Scr (mg/dL)** | 0.65 ± 0.03 | 0.91 ± 0.12 | 0.97 ± 0.11 | 1.17 ± 0.15* | 0.82 ± 0.06 | 0.83 ± 0.06 |
| **Ucr (mg/dL)** | 65 ± 8 | 21 ± 4* | 24 ± 3* | 25 ± 4* | 30 ± 5* | 26 ± 2* |
| **CrCl (mg/min/RBSA)** | 31 ± 3 | 20 ± 6 | 23 ± 5 | 16 ± 2* | 35 ± 10 | 24 ± 2 |
| **BUN (mg/dL)** | 51 ± 4 | 105 ± 13* | 88 ± 8* | 101 ± 15* | 84 ± 7* | 82 ± 4* |
| **Renal Hypertrophy** | 3.4 ± 0.1 | 5.0 ± 0.3* | 5.2 ± 0.2* | 4.9 ± 0.5* | 5.3 ± 0.4* | 4.6 ± 0.1* |

